# Metagenomic 16S rRNA gene sequencing survey of *Borrelia* species in Irish samples of *Ixodes ricinus* ticks

**DOI:** 10.1101/496950

**Authors:** John S Lambert, Michael John Cook, John Eoin Healy, Ross Murtagh, Gordana Avramovic, Sin Hang Lee

## Abstract

Lyme borreliosis is a systemic infection caused by tick-borne pathogenic borreliae of the *Borrelia burgdorferi* sensu lato complex or of the more heterogeneous relapsing fever borrelia group. Clinical distinction of the infections due to different borrelia species is difficult. Accurate knowledge of the prevalence and the species of borreliae in the infected ticks in the endemic areas is valuable for formulating appropriate guidelines for proper management of this infectious disease. The purpose of this research was to design a readily implementable protocol to detect the divergent species of borreliae known to exist in Europe, using Irish samples of *Ixodes ricinus* ticks as the subject for study. Questing *I. ricinus* nymph samples were taken at six localities within Ireland. The crude DNA of each dried tick was extracted by hot NH_4_OH and used to initiate a same-nested PCR with a pair of borrelial genus-specific primers to amplify a highly conserved 357/358 bp segment of the 16S rRNA gene for detection and as the template for Sanger sequencing. To distinguish *B. garinii* from *B. burgdorferi* and to discriminate the various strains of *B. garinii*, a second 282 bp segment of the 16S rRNA gene was amplified for Sanger sequencing. A signature segment of the DNA sequence excised from the computer-generated electropherogram was submitted to the GenBank for BLAST alignment analysis. A 100% ID match with the unique reference sequence in the GenBank was required for the molecular diagnosis of the borrelial species or strain. We found the overall rate of borrelial infection in the Irish tick population to be 5%, with a range from 2% to 12% depending on the locations of tick collection. At least 3 species, namely *B. garinii, B. valaisiana* and *B. miyamotoi*, are infecting the ticks collected in Ireland. The isolates of *B. garinii* were confirmed to be strain BgVir, strain Bernie or strain T25. Since antigens for diagnostic serology tests may be species- or even strain-specific, expanded surveillance of the species and strains of the borreliae among human-biting ticks in Ireland is needed to ensure that the antigens used for the serology tests do contain the epitopes matching the antibodies elicited by the borrelial species and strains in the ticks cohabitating in the same environment.

## Introduction

Lyme borreliosis (or Lyme and related borreliosis) can be caused by many members of the *Borrelia burgdorferi* sensu lato (B.b.s.l) complex [1] and the more heterogeneous group of relapsing fever borreliae [2] which are transmitted to humans through the bite of infected ticks of the *Ixodes* genus, including *I. ricinus, I. scapularis, I. pacificus* and *I. persulcatus*. In Europe, the pathogenic B.b.s.l. species include *Borrelia afzelii, B. garinii, B. bavariensis, B. spielmanii, B. valaisiana, B. lusitaniae* and rarely *B. burgdorferi* [1, 3-6]; the major hard tick relapsing fever borrelia is *B. miyamotoi* [7].

Lyme borreliosis at its early stage of infection can usually be effectively treated with timely, appropriate antibiotics to prevent deep tissue damage along with the associated clinical manifestations resulting from host immune response to various spirochetal products or components. However, if not treated early, within days to weeks, the borrelial spirochetes disseminate from the site of the tick bite to other regions of the body [8].

Since Lyme borreliosis only occurs in the endemic areas where the tick population is infected with pathogenic borrelial species, accurate surveys of the borrelial infections of the ticks collected in the potential endemic areas can provide valuable information on the possible existence of any of these Lyme and related borrelioses, and can serve as guidelines for selection of the proper antigens for diagnostic serology tests which may be species- or even strain-specific [9-11].

In the past, the methods used for borrelial surveys were either designed to detect *B. burgdorferi* sensu lato [1] or to detect *B. miyamotoi* [7, 12] in ticks; the PCR primers used for these metagenomic assays were unable to amplify a conserved segment of DNA shared by all species of borreliae. Only one study used a pair of general PCR primers which can amplify a segment of 16S rRNA gene of both *B. burgdorferi* sensu lato and *B. miyamotoi* for real-time PCR screening [13]. But the interprimer DNA segment of the latter real-time PCR product is only 25 bp long which is too short for automated Sanger sequencing that is required to confirm the molecular diagnosis of *B. miyamotoi* [7, 12, 13]. In addition, the dye-labeled probe 6FAM-TTCGGTACTAACTTTTAGTTAA used for the real-time PCR screening [13] may miss the species of *B. valaisiana* and *B. lusitaniae* whose corresponding complementary sequences in this segment are TTAACTGAAAGTTAGTACCGAA (Sequence ID: NR_036807) and TTAACTAACAGTTAGTACCGAA (Sequence ID: AB091822), respectively, not fully matched with the sequence of the probe designed for other species of the *B. burgdorferi* sensu lato complex.

In the current study, we used a pair of genus-specific PCR primers to amplify a highly conserved segment with single-nucleotide polymorphisms of the borrelial 16S rRNA gene shared by all known pathogenic borrelial strains to survey the borrelial infections among the *I. ricinus* ticks collected in Ireland. Since this PCR amplicon is 357/358 bp long, the PCR products can be used as the template for direct Sanger sequencing for amplicon validation and for speciation [14]. This metagenomic 16S rRNA gene sequencing assay is most suitable for molecular diagnosis of borrelial infections in human-biting ticks and in clinical specimens in Europe because of the great diversity of causative agents in European Lyme borreliosis which needs a broad-spectrum tool to detect the target DNA from various borrelial strains and to prepare the template for Sanger sequencing to ensure diagnostic accuracy.

The value of routine metagenomic 16S rRNA gene sequencing is well recognized for detecting the presence of pathogenic bacteria in the environment and in clinical specimens because single nucleotide polymorphisms in the 16S rRNA gene may discriminate closely related bacterial species or indicate a mutant expressing different patterns of synthesized proteins with potential implication in antibody epitopes, virulence and drug resistance of the bacteria [15-19]. However, the usefulness of 16S rRNA gene sequencing as a tool in microbial identification is dependent upon deposition of the complete unambiguous 16S rRNA gene nucleotide sequences into a public database, such as that in the GenBank and applying the correct base labels to each sequence detected for alignment analysis [20]. Indeed, the validity of the results of studying borrelia distribution in *I. scapularis* ticks using PCR and DNA sequencing has been questioned because the DNA sequences claimed to have been detected were questionable [21]. Good-quality sequence traces in Sanger sequencing are essential for molecular diagnosis and typing of borreliae [5] in the era of precision medicine.

In the present study, we aimed to develop a protocol for using a single pair of genus-specific PCR primers to amplify a highly conserved segment with hypervariable regions of the borrelial 16S rRNA gene for detection of all species of *Borrelia* infecting the *I. ricinus* ticks collected in Ireland and to use the positive crude nested PCR products as the templates for direct Sanger sequencing to determine the species of the borrelia detected.

## Materials and Methods

Unfed, questing *I. ricinus* nymphs were collected by “flagging” which involves brushing the vegetation with a white towel from which the ticks can then be removed. In late May and early June 2018, samples were taken at six localities within Ireland, designed to provide a representative view of tick borrelial infection across the country. The following areas were sampled (county in parenthesis): Killarney (Kerry), Kilmacthomas (Waterford), Clifden (Galway-West), Portumna (Galway-South), Glendalough (Wicklow), and Glenveagh (Donegal) and their locations are shown on the accompanying map of Ireland (Fig 1).

**Fig 1.**
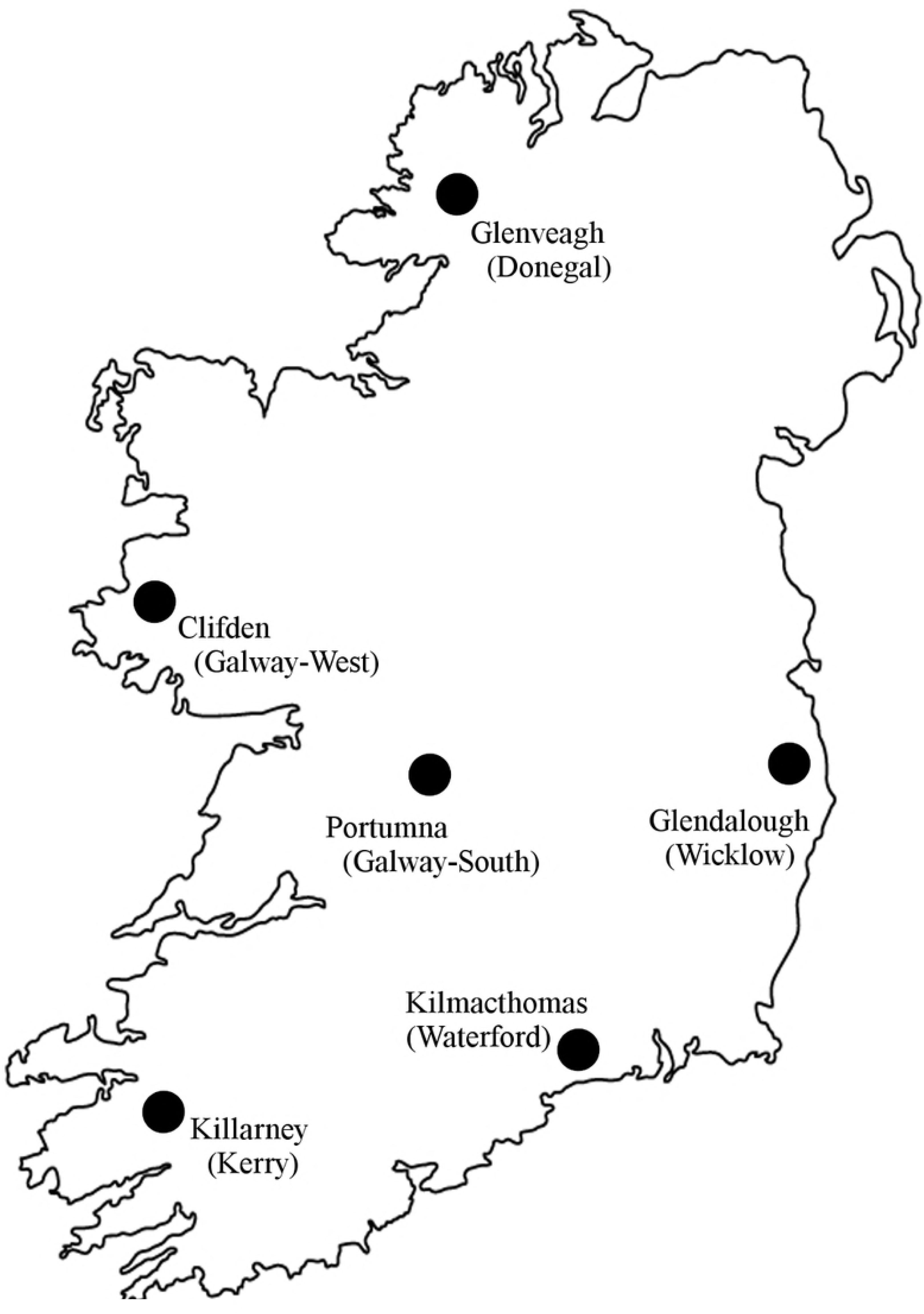
Sampling locations in Ireland for *Ixodes ricinus* ticks in May & June 2018.

Each individual sample was made up of at least 5 sub-samples taken at different points within a locality to minimise any sampling bias. Distance between sub-sampling points was never less than 100 m.

The collected ticks were disinfected in 70% ethanol, air-dried on filter paper and sent to Milford Molecular Diagnostics Laboratory in Milford, Connecticut, U.S.A. to be tested. The general procedure for extraction of the crude DNA from archived ticks and for Sanger sequencing detection of the borrelial 16S rRNA gene previously published [22] was followed. Initially, 300 ticks were analysed, 50 from each of the six sampled locations listed above.

On the day of testing, each dried tick was placed in a 1.5 mL plastic tube and immersed in 300 μL of 0.7 mol/L NH_4_OH overnight at room temperature. On the following day, the test tubes were heated at 95°C to 98°C for 20 minutes with closed caps, followed by 10 minutes with open caps. After the test tubes were cooled to room temperature and the carcass of the tick was discarded, 700 μL of 95% ethanol and 30 μL of 3 mol/L sodium acetate (Sigma) were added to each NH_4_OH digestate. The precipitated crude DNA was spun down in the pellet after centrifugation at ∼ 16,000 x g for 5 minutes, washed in 1 mL of 70% ethanol, air dried, and re-dissolved in 100 μL of tris(hydroxymethyl)aminomethane hydrochloride–EDTA (TE) buffer, pH 7.4 (Sigma) by heating the DNA extract at 95°C to 98°C for 5 minutes. After centrifugation, 3µL of the crude DNA extract in the supernatant was used to initiate a primary PCR, followed by a same-nested PCR using a pair of M1/M2 borrelial genus-specific primers in a total 25 μL low-temperature PCR mixture for a 30-cycle amplification at primary PCR, followed by another 30-cycle amplification at the same-nested PCR. In carrying out the same-nested PCR, a single pair of M1 (5′-ACGATGCACACTTGGTGTTAA-3′) and M2 (5′ TCCGACTTATCACCGGCAGTC-3′) primers were used for both primary PCR and nested PCR so that a small amount of the primary PCR products was re-amplified with the same pair of PCR primers in a new PCR mixture [14]. An original target DNA segment in the PCR mixture might have been amplified for 60 cycles exponentially by the same pair of primers to increase the sensitivity of detection. The PCR amplicons of the 357-bp 16S rRNA gene segment of the *B. burgdorferi* sensu lato complex and the corresponding 358-bp 16S rRNA gene segment of the relapsing fever borreliae, both defined by the M1 and M2 PCR primer pair, were visualized by agarose gel electrophoresis of the nested PCR products. The same-nested PCR products were used as the template for Sanger reaction without purification.

For DNA sequencing, the positive nested PCR products were transferred by a micro-glass rod into a Sanger reaction tube containing 1 μL of 10 μmolar sequencing primer, 1 μL of the BigDye^®^ Terminator (v 1.1/Sequencing Standard Kit), 3.5 μL 5× buffer, and 14.5 μL water in a total volume of 20 μL for 20 enzymatic primer extension/termination reaction cycles according to the protocol supplied by the manufacturer (Applied Biosystems, Foster City, CA, USA). After a dye-terminator cleanup with a Centri-Sep column (Princeton Separations, Adelphia, NJ, USA), the reaction mixture was loaded in an automated ABI 3130 four-capillary Genetic Analyzer for sequence analysis [14].

Sanger sequencing of positive 357/358 bp M1/M2 same-nested PCR products is capable of accurate identification of many species including, *B. valaisiana, B. afzelii, B. mayonii, B. spielmanii, B. lusitaniae, B. recurrentis, B. miyamotoi, B. hermsii, B. lonestari, B. coriaceae* and several other members in the relapsing fever group based on known species-specific single-nucleotide polymorphisms in the gene segment [14]. However, DNA sequencing of this 16S rRNA gene segment cannot distinguish *B. garinii* and *B. bavariensis* from *B. burgdorferi* sensu stricto or discriminate the heterogeneous strains within the species of *B. garinii* because their base sequences defined by the M1 and M2 PCR primers are identical.

Ireland is an island state geographically at the periphery of Europe. It is important at least for the current study to accurately determine the species or strains of the *B. burgdorferi* sensu lato detected in the ticks because the causative agents of Lyme borreliosis are highly heterogeneous in Europe. [1, 3-6] In North America, species differentiation among members of the *B. burgdorferi* sensu lato complex was initially considered unnecessary in a routine diagnostic laboratory because practically all *B. burgdorferi* sensu lato isolates detected in ticks and in clinical specimens in the United States were presumed to be *B. burgdorferi* sensu stricto [23]. Nucleic acid-based diagnostic tests had been designed to detect *B. burgdorferi* sensu stricto only [24]. However, it is now recognized that the borrelia strains causing clinical Lyme disease in the United States are in fact quite diverse, especially after *B. miyamotoi* infection in humans was reported in 2013 [25]. According to the current *Viewpoints* of a group of Lyme disease experts, a “core genome” shared by all isolates capable of causing clinical Lyme borreliosis is needed for direct detection diagnosis even in the United States [26].

The “genus-specific” M1/M2 PCR primer pair can amplify a “core genome” of all pathogenic borreliae for the purpose of detection. However, to design a pair of reliable PCR primers to amplify a segment of borrelial 16S rRNA gene with single-nucleotide polymorphisms among various borrelial strains for further species or strain differentiation without co-amplification of the unwanted DNA extracted from other bacteria also present in the ticks turned out to be challenging. In molecular diagnosis, the size of the PCR amplicons for detecting a small quantity of bacterial target DNA in a pool of non-target bacterial DNAs are usually below 300 bp in size [27-30] to avoid loss of sensitivity. It took several weeks of experimental work before we found 3 PCR primers to generate a 282-bp heminested PCR amplicon useful as the template for Sanger sequencing to distinguish *B. garinii* from *B. burgdorferi* and to discriminate among the various *B garinii* strains. The sequences of these 3 heminested PCR primers are listed as follows.

Primary PCR Forward Primer Bg1: 5’-GACGTTAATTTATGAATAAGC −3’

Primary PCR Reverse Primer Bg 6: 5’-TTAACACCAAGTGTGCATCGT – 3’

Heminested PCR Forward Primer Bg5: 5’-CGGGATTATTGGGCGTAAAGGGTGAG-3’

Heminested PCR Reverse Primer Bg 6: 5’-TTAACACCAAGTGTGCATCGT – 3’

Based on the reference sequences retrieved from the GenBank, the Bg5/Bg6 heminested PCR primer pair defines a 282 bp segment of the borrelial 16S rRNA gene with single-nucleotide polymorphisms which can be used to discriminate *Borrelia burgdorferi* strain B31 (ID# CP019767), *Borrelia garinii* BgVir (ID# CP003151), *Borrelia garinii* strain Bernie (ID# D89900), *Borrelia garinii* strain T25 (ID# AB035388) and *Borrelia garinii* strain L20 (ID# X85198) for the purpose of routine molecular diagnosis.

However, after the first 300 ticks were tested, it was realized that the free DNA of the borrelia 16S rRNA gene in the crude DNA extract from the ticks was not stable on storage even at −°C. By the time when the Bg5 and Bg6 heminested PCR primers were readied to be put into routine practice, the borrelial 16S rRNA gene DNA in 9 of the 12 samples initially found to be positive for *B. burgdorferi* sensu lato already degraded and were no longer amplifiable with any PCR primers. Therefore, an additional series of 50 ticks from each of the Portumna and Kilmacthomas samples were analysed for the specific purpose of species confirmation and strain determination of the *B. garinii* isolates.

## Results

### Multiple borrelial species found in *I. ricinus* ticks in Ireland

The same-nested PCR amplification of a 357/358 bp segment of borrelial 16S rRNA gene by the M1/M2 genus-specific PCR primers followed by Sanger sequencing of the nested PCR products [14] provided metagenomic evidence of *B. burgdorferi* sensu lato (B.b.s.l.), *B. valaisiana* and *B. miyamotoi* infection in the ticks collected in Ireland. Samples of the 16S rRNA gene sequencing with the M2 primer in support of these molecular diagnoses are illustrated by the 3 selected electropherograms presented in Fig 2.

**Fig 2.**
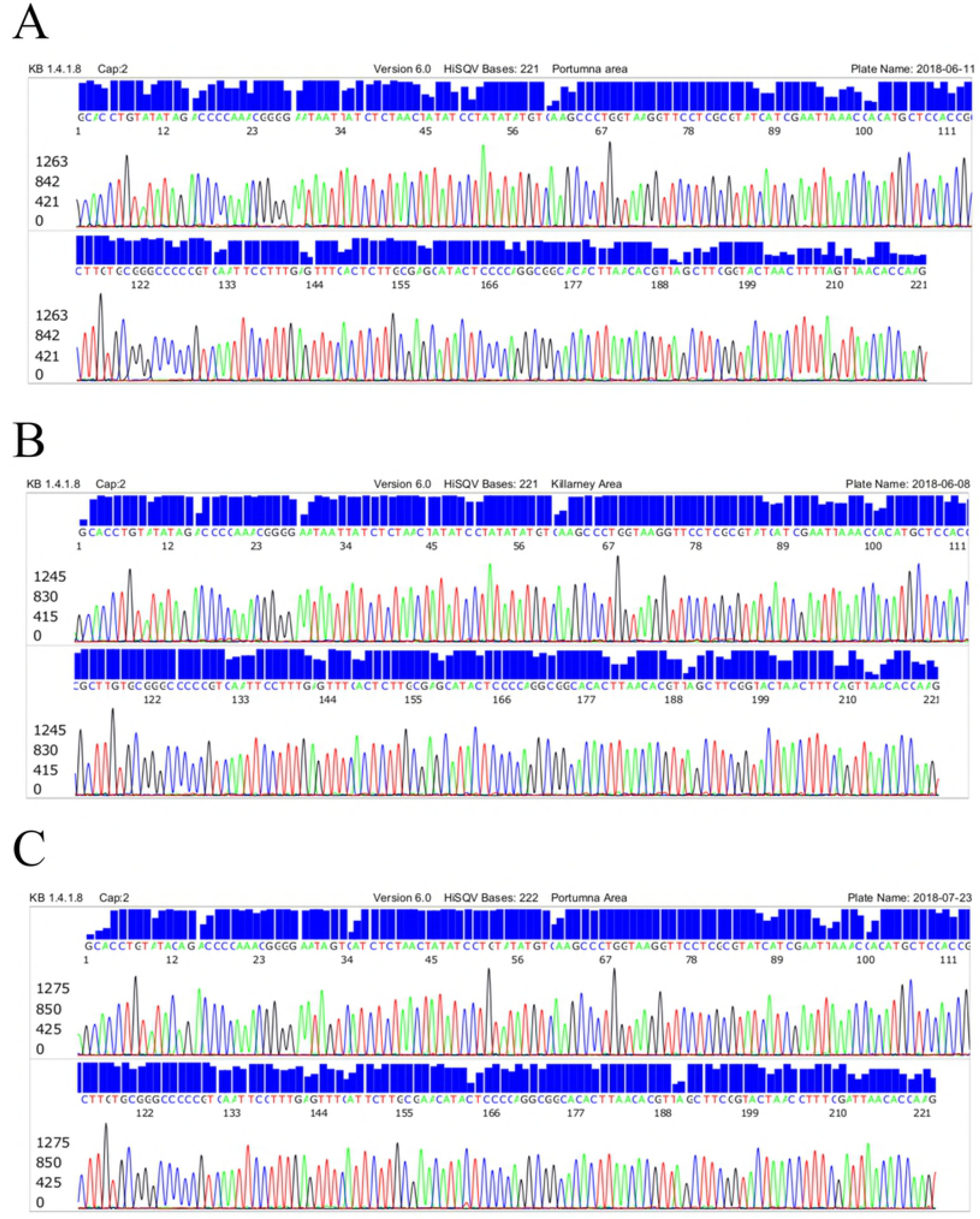
Samples of computer-generated electropherograms of 16S rRNA gene sequences representing the 3 borrelial groups detected in the ticks collected in Ireland. (A) 221 bases Portumna area – *B. burgdorferi* sensu lato (GenBank ID# CP019767). (B) 221 bases Killamey Area – *B. valaisiana* (GenBank ID# AB091815). (C) 222 bases Portumna Area – *B. miyamotoi* (GenBank ID# KM007554).

### Strain diversity of *B. garinii*

The *B. burgdorferi* sensu lato isolates other than *B. valaisiana, B. spielmanii, B. mayonii* or *B. lusitaniae* cannot be speciated by the DNA sequence illustrated in Fig 2A, based on single-nucleotide polymorphisms of the 16S rRNA gene DNA sequences retrieved from the GenBank [14]. Sanger sequencing of the PCR amplicon with the reverse M1 primer proved that none of the *B. burgdorferi* sensu lato isolates were *B. afzelii*. A 177-base sequence of the 282-bp amplicon defined by the Bg5 and Bg6 heminested PCR primers distinguished the heterogeneous strains of *B. garinii* from *B. burgdorferi* and *B. bavariensis* due to the presence of single-nucleotide polymorphisms among strain BgVir {1}, strain 25 {2} and strain Bernie {3} of the *B. garinii* species and distinguished these strains from *B. burgdorferi* sensu stricto {4}. Alignment of these four 177-base reference sequences retrieved from the GenBank showing single-nucleotide polymorphisms is presented below with the ending 26-base G5 primer site underlined.

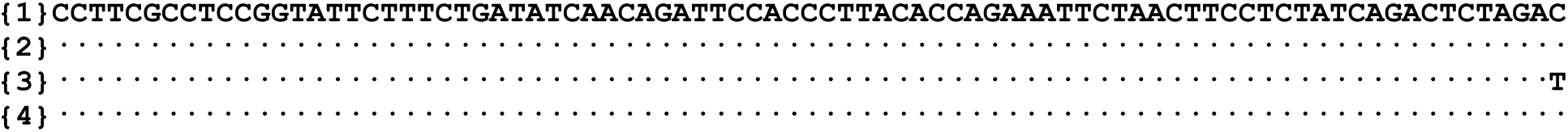

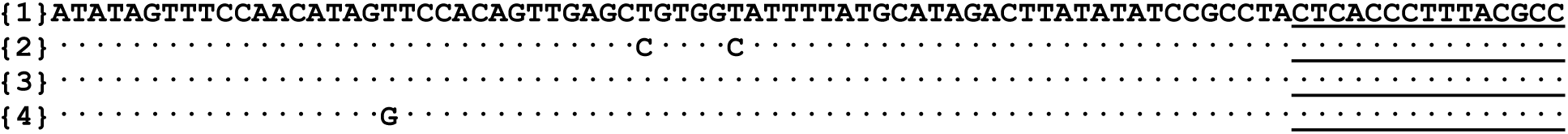

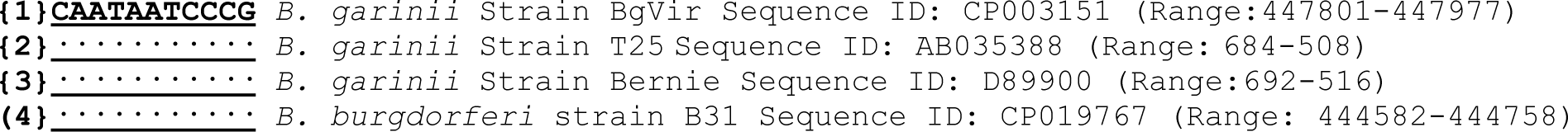

The corresponding electropherograms of these four 177-base sequences (1-4), using the Bg6 sequencing primer, are illustrated in Fig 3, arranged from top to bottom in numerical order of sequences 1-4 as Strain BgVir, Strain T25, Strain Bernie and *B. burgdorferi* strain B31.

**Fig 3.**
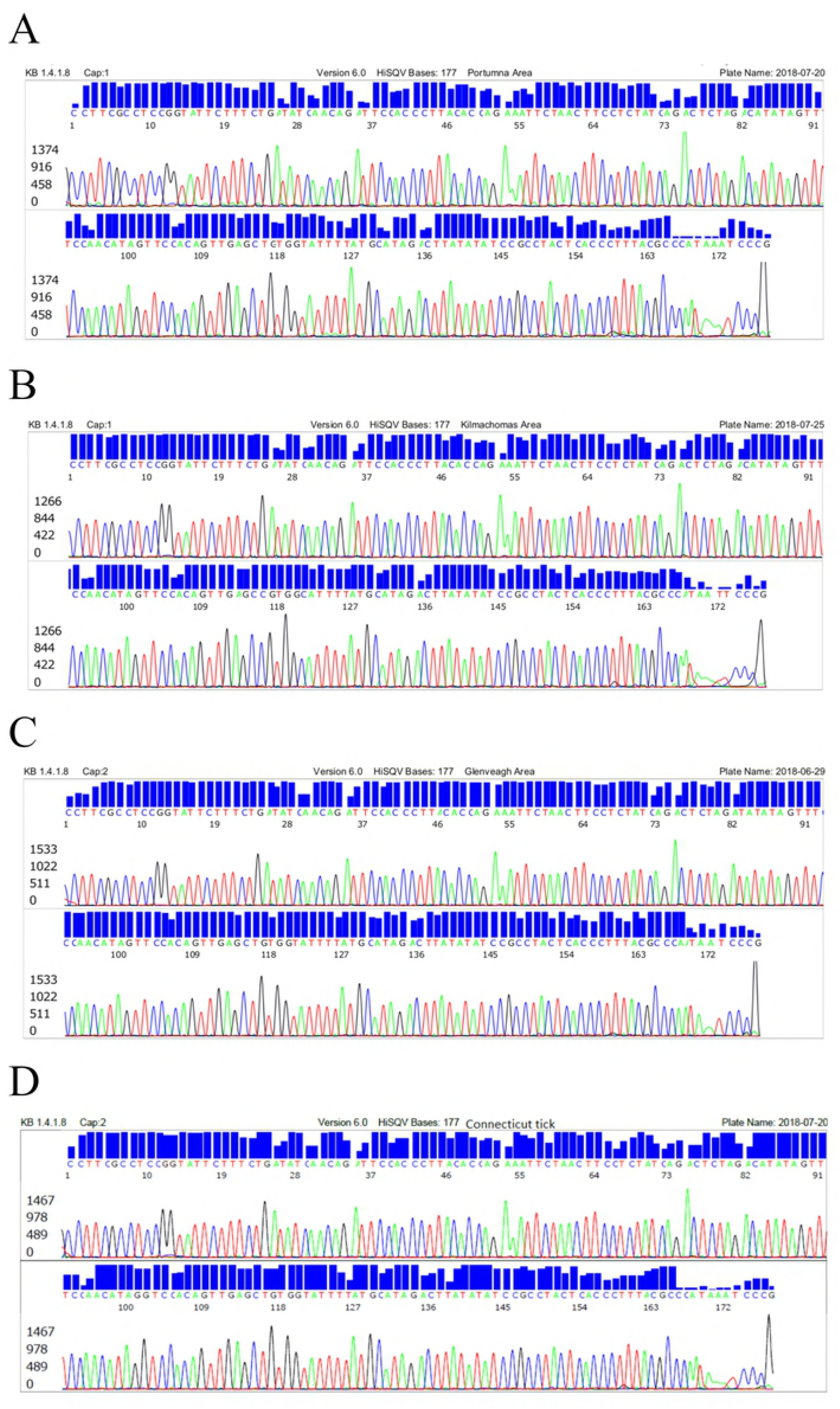
DNA sequencing electropherograms for distinguishing *B. garinii strains*. (A) Strain BgVir. (B) Strain T25. (C) Strain Bernie. (D) *B. burgdorferi* strain B31.

A reverse sequencing with Bg5 primer of the 282-bp PCR amplicon confirmed that the isolates identified as strains BgVir were not strain SL20 (Sequence ID: X85198) because there is no single-nucleotide polymorphism located near the Bg6 primer site of the amplicon which is unique for strain SL20.

### Tick infection rates varied with locations

As stated above, we conducted two surveys. Survey 1 consisted of 300 ticks collected in six locations showing 15 of the 300 ticks (5%) infected with *B. burgdorferi* sensu lato of which 3 were confirmed to be *B. garinii* (2 being strain Bernie and 1 strain BgVir), 2 were *B. valaisiana* and 10 were not speciated. Survey 2 consisted of 100 ticks collected in two of the six locations previously surveyed, showing 5 of the 100 ticks (5%) infected by borrelial species, including 4 isolates of *B. garinii* (3 being strain BgVir and 1 being strain T25) and 1 isolate of *B. miyamotoi*. The borrelial infection rates varied from 2% to 12 % depending on the locations of the tick collection. It is noteworthy that all *B. burgdorferi* sensu lato isolates in Survey 2 were confirmed to be *B. garinii* as the Bg5/BG6 PCR followed by Sanger sequencing for speciation was carried out without delay after initial detection by M1/M2 PCR. The final species distributions in different locations were summarized in Table 1.

**Table 1.** Incidence of *Borrelia* species in Irish *Ixodes ricinus* tick samples (Nymphs).

| Location | Sample size | No. Positive | % Positive | CI 95% | *Borrelia* species |
| --- | --- | --- | --- | --- | --- |
| Killarney (Kerry) | 50 | 1 | 2% | (0% - 6%) | *B. valaisiana* |
| Killmacthomas (Waterford) | 50 | 3 | 6% | (0% - 13%) | *B.b s.l* |
| Killmacthomas (Survey 2) | 50 | 2 | 4% | (0% - 9%) | 1x *B. garinii* strain T25<br>1x *B. garinii* strain BgVir |
| Portumna (Galway-South) | 50 | 6 | 12% | (3% - 21%) | *B.b s.l* |
| Portumna (Survey 2) | 50 | 3 | 6% | (0% - 13%) | 1x *B. miyamotoi*<br>2x *B. garinii* strain BgVir |
| Gelendalough (Wicklow) | 50 | 1 | 2% | (0% - 6%) | *B. valaisiana* |
| Glenveagh (Donegal) | 50 | 3 | 6% | (0% - 13%) | 1x *B. garinii* strain BgVir<br>2x *B. garinii* strain Bernie |
| Clifden (Galway-West) | 50 | 1 | 2% | (0% - 6%) | *B.b s.l* |
| Total | 400 | 20 | 5% | (3% - 7%) |  |
| Of 20 infected ticks |  |  |  |  |  |
| *B. garinii* |  |  |  | 35% | (7/20) |
| *B. miyamotoi* |  |  |  | 5% | (1/20) |
| *B. valaisiana* |  |  |  | 10% | (2/20) |
| Percent other *B.b s.l* not speciated |  |  |  | 50% | (10/20) |

*B.b s.l* = *Borrelia burgdorferi* sensu lato.
CI 95% = Confidence Interval for 95% probability.

#### Selection of PCR primers for 16S rRNA gene PCR and sequencing

For routine metagenomic DNA sequencing diagnosis of borrelial infections, it would be ideal to use one single pair of broad-spectrum PCR primers which can amplify a highly conserved segment of the 16S rRNA gene of all species in the *Borrelia* genus and also can discriminate between different borrelial species, and does not amplify the DNAs of other environmental bacteria which may co-exist in the sample, in a nested PCR setting to prepare templates for DNA sequencing. However, such an ideal specific PCR primer pair is difficult to design. We use the “genus-specific” M1/M2 primer pair to generate a PCR amplicon for Sanger reaction to detect all species of the B.b.s.l. complex and the relapsing fever borreliae, in particular *B. miyamotoi*. But the amplicons generated by the M1/M2 PCR primers do not contain the sequence polymorphisms for discriminating between certain species in the B.b.s.l. complex and between certain species of the relapsing fever borreliae. Hence, we chose a heminested PCR system to amplify an adjacent 282 bp segment defined by the Bg5 and Bg6 primers which is known to have single-nucleotide polymorphisms to prepare the template for further species differentiation by DNA sequencing (Fig 3). However, when the Bg5/Bg6 nested PCR products were used as the DNA sequencing template for confirmation of *B. miyamotoi* in tick samples, the electropherograms showed numerous ambiguous base calling peaks as a result of co-amplification of unwanted DNAs in the sample extract (Fig 4) whereas the sequencing electropherogram generated by the M1/M2 primer PCR products from the same tick extract showed no ambiguous base calling labels (Fig 2C).

**Fig 4.**
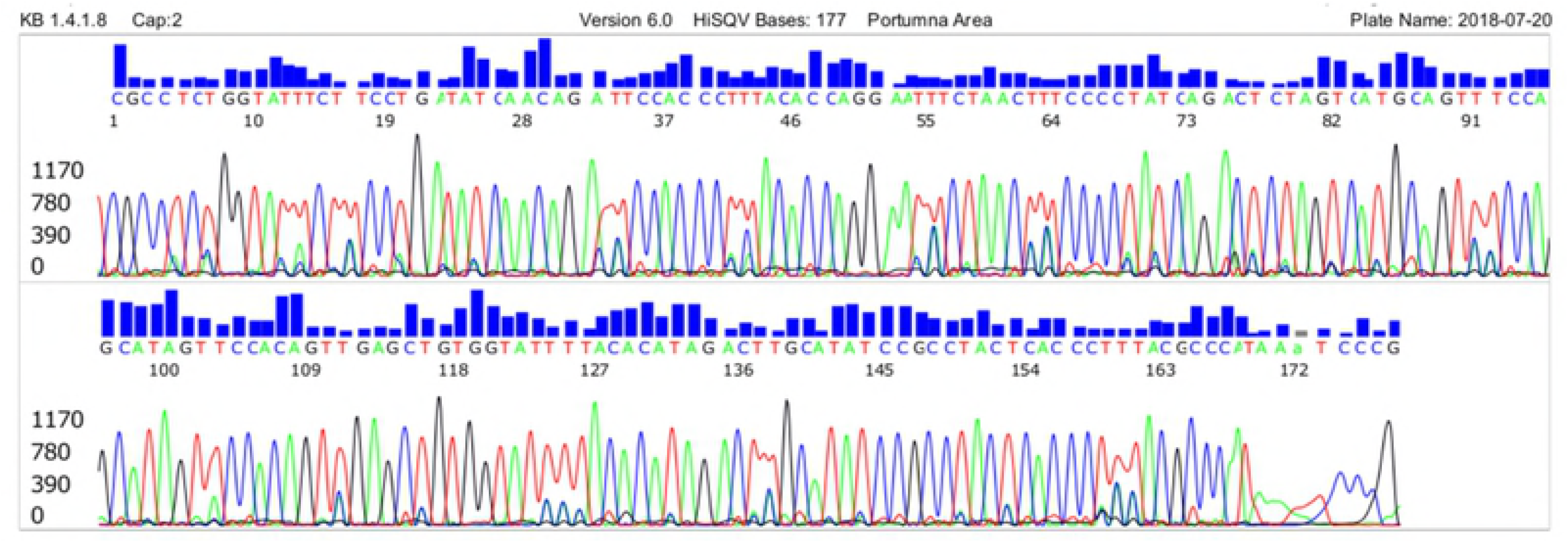
DNA sequencing of Bg5/Bg6 primer nested PCR products of *B. miyamotoi* 16S rRNA gene showing many ambiguous bases, compared to sequencing of M1/M2 primer nested PCR products of the same sample (see Fig 2C)

It is well known that in using metagenomic 16S rRNA gene sequencing for target bacterial identification in samples containing a mixture of bacteria, the selected target variable regions of the 16S rRNA gene defined by different PCR primers have a major impact on the analysis results [31, 32].

### Degradation of 16S rRNA gene DNA in samples

We also observed that free borrelial 16S rRNA gene DNA in crude extracts from the ticks was unstable even in TE buffer stored at −20°C for an extended period of time. For example, when the NH_4_OH extracts of the 50 ticks from the Portumna area were processed for M1/M2 same-nested PCR screening on July 20, 2018, 3 samples were positive for borrelial 16S rRNA gene DNA, with a robust 357/358 bp band on lanes 11, 15 and 25 (Fig 5) which eventually proved to be *B. garinii* strain BgVir, *B. miyamotoi* and *B. garinii* BgVir, respectively. When the NH_4_OH extracts of the 50 ticks from the Kilmacthomas area were processed for M1/M2 same-nested PCR screening on July 24, 2018, 2 samples were positive for borrelial 16S rRNA gene DNA, with a robust 357/358 bp band on lanes 372 and 382 (Fig 5) which eventually proved to be *B. garinii* strain BgVir and *B. garinii* strain T25, respectively. However, when the same-nested PCR was repeated on these 5 NH_4_OH extracts after 7 days and 3 days storage in a −20°C freezer, respectively, the 16S rRNA gene DNA in sample 11 was no longer detectable and the intensities of the nested PCR bands using the same extracts of samples 15, 25, 372 and 382 as primary PCR templates for amplification under identical experimental conditions decreased markedly over a period of 3-7 days, as demonstrated on the agarose gel dated July 27, 2018 (Fig 5). The image of the gel electrophoresis dated July 27, 2018 also showed that nested PCR is generally required for detection of borrelial infections by 16S rRNA gene analysis. Primary PCR products are usually invisible after gel electrophoresis.

**Fig 5.**
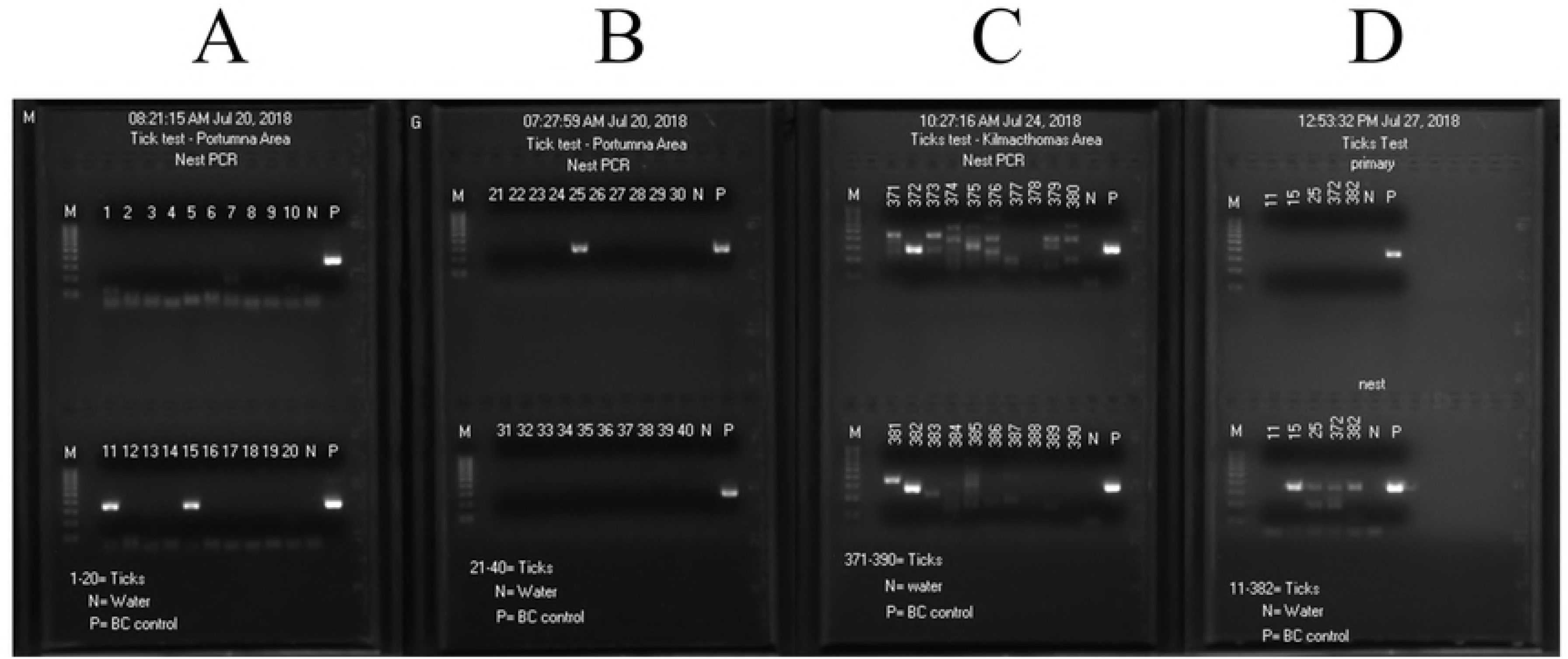
Images of agarose gel electrophoresis. Images on July 20 and July 24, 2018 (A-C) show positive 357/358 bp borrelial 16S rRNA gene nested PCR amplicons in five positive tick samples on the day of DNA extraction (Lanes 11, 15, 25, 372 and 382). After storage of the DNA extracts at −20°C, the same DNA preparations were used to repeat primary and nested PCR on July 27, 2018 (D), showing no nested PCR products for sample 11; the amount of nested PCR products on samples 15, 25, 372 and 382 was significantly reduced. N=negative control; P= positive *Borrelia coriaceae* control. M=molecular ruler. Note numerous non-specific PCR amplifications on panel for samples 371-390.

## Discussion

DNA sequencing of the 16S ribosomal RNA gene is a well-established method to compare and identify bacteria [34]. Sequence analysis of the 16S rRNA gene is a reliable method for species determination of the *B. burgdorferi* sensu lato complex and the relapsing fever borreliae [34-38].

Routine metagenomic 16S rRNA gene sequencing test for spirochetemia was initially developed for the diagnosis of human Lyme disease in the United States [22, 39] where all isolates of pathogenic *B. burgdorferi* sensu lato detected were presumed to be *B. burgdorferi* sensu stricto [23]. Subsequently, a pair of broad-spectrum genus-specific M1/M2 primers was introduced to amplify a highly conserved 357/358 bp segment of the borrelial 16S rRNA gene as the template for Sanger sequencing to include *B. miyamotoi* as the target for detection [14] after *B. miyamotoi* was found in ticks [13] and in a patient [25]. In the era of precision medicine, microbiological diagnosis of bloodstream infections, such as borrelial spirochetemia, may rely on using accurate bacterial nucleic acid analysis to establish if blood culture is not an option [40]. Base calling electropherograms generated in the automated Sanger sequencing process as those shown in Fig 2 and 3 may offer objective physical evidence for visual analysis in all nucleic acid-based diagnostic tests for infectious diseases, as proposed by some bacteriologists who are seriously concerned about the accuracies of molecular diagnosis for health care improvement [41,42].

Our study shows that *I. ricinus* ticks in Ireland are infected by a diversity of pathogenic borreliae. The degree of species and strain diversity may prove to be greater if the number of ticks being surveyed is increased. For example, in our two surveys *B. miyamotoi* and *B. garinii* strain T25 were demonstrated only during Survey 2, but not in Survey 1 (Table 1). Importantly, our results demonstrate that *Borrelia*-infected tick populations exist in the east and south-east of Ireland, areas hitherto not considered to be significantly tick-infested and hence not areas of risk to humans for contraction of Lyme borreliosis.

Historically, the only surveys of *B. burgdorferi* sensu lato in ticks collected in Ireland were carried out more than 20 years ago, using reverse line blot (RLB) for detection and reporting infection rates to be between 3.5% and 26.7% depending on the locations of tick collection at that time [43, 44]. In the current study, we used metagenomic 16S rRNA gene sequencing to survey the *I. ricinus* ticks collected from the wooded areas in six counties of Ireland and found the overall rate of borrelial infection to be 5%, with a range from 2% to 12% depending on the locations of tick collection (Table 1). Comparing our results with those published 20 years ago is difficult because there is a significant difference between the RLB and Sanger sequencing technologies in DNA test accuracy. Nevertheless, we found that 6% of the nymphs collected in Glenveagh were infected by two different strains of *B. garinii*, compared to 3.6% (5/141) infected of the nymphs collected in Glenveagh National Park which were reported to be positive for *B. valasiana* (labeled as strain VS116) 20 years earlier [44]. As DNA dot-hybridization assay is prone to false positive results when no DNA sequencing confirmation is performed for validation [45], it is not possible to determine whether the local strains of *B. valaisiana* in the ticks in Glenveagh have been replaced by *B. garinii* over the past 20 years. As reported here, with Sanger sequencing it is beyond reasonable doubt that at least 3 pathogenic borrelial species, namely *B. garinii, B. valaisiana* and *B. miyamotoi*, are infecting the ticks collected in Ireland. The isolates of *B. garinii* are strain BgVir strain, strain Bernie or strain T25. This is the first time that *B. miyamotoi* was detected in in a tick collected in Ireland although it was previously reported in *I. ricinus* ticks in Britain [46].

Extracellular bacterial 16S rRNA gene DNA is prone to degradation in the presence of environmental substances [47], a phenomenon which was recognized when borrelial 16S rRNA gene DNA in human clinical specimens was studied [48] and was also observed in the current study on borrelial DNA extracts from infected ticks (Fig 5). However, this kind of DNA degradation seems to be not inevitable. For example, as shown in Table 1, all of the 3 positive extracts of the ticks collected from Glenveagh in Survey 1 contained adequate preserved 16S rRNA gene DNA to be further speciated as *B. garinii* after initially being grouped under the *B. burgdorferi* sensu lato complex whereas all the extracts from the ticks collected from other locations lost their amplifiable 16S rRNA gene DNA under identical storage conditions. For comparison, all four extracts of the ticks in Survey 2 found to be positive for *B. burgdorferi* sensu lato contained adequate 16S rRNA gene DNA for speciation which confirmed that the isolates initially diagnosed as *B. burgdorferi* sensu lato were in fact strains of *B. garinii* when the speciation PCR was carried out within 24 hours. To avoid DNA degradation when using metagenomic sequencing of borrelial 16S rRNA gene for molecular diagnosis, PCR amplification should be carried out without delay after NH_4_OH extraction.

For the purpose of patient care, it is probably not necessary to determine the species or strain of the borrelia detected in the blood sample of a patient suffering from acute *B. burgdorferi* sensu lato or *B. miyamotoi* infection confirmed by DNA sequencing in order to initiate timely antibiotic treatment. But, for serological diagnosis a knowledge of the borrelial species and strains carried by the human-biting ticks collected in the endemic areas is crucial since polymorphism of ospC [9-11] and variation of the VlsE antigens among species and strains of *B. burgdorferi* sensu lato [49] have been well documented in the literature. There is limited information regarding the sensitivity of commercial tests for different species of borrelia. The manufacturers of Western Blot antibody tests specify the target species. In Europe these are typically *B. afzelii, B. burgdorferi* and *B. garinii*. A study of the most frequently used C6 synthetic peptide ELISA test for initial screening does appear to have variable sensitivity for some species [50] but with no data available for many of the European species including *B. valaisiana* identified as prevalent in Ireland in this study.

In summary, our study confirms that the genus-specific M1/M2 PCR primers can amplify a highly conserved segment of the borrelial 16S rRNA gene for Sanger sequencing-based molecular diagnosis of tick-borne borreliae. Three species of *borrelia* were identified with *B. garinii* the most common (60% of those speciated), followed by *B. valaisiana* (20%), and for the first time in Ireland *B. miyamotoi* (10%) has been identified. *Borrelia* DNA was identified in 12% of the ticks collected in the Portumna area from the first survey, and in up to 6% of the ticks in Glenveagh and Kilmathomas areas. Expanded surveillance of the species and strains of the borreliae among human-biting ticks in Ireland is needed to ensure that the antigens used for the serology tests do contain the epitopes matching the antibodies elicited by the borrelial species and strains in the ticks cohabitating in the same environment.

## Author Contributions

**Conceptualization:** John S. Lambert, John Eoin Healy.

**Data Curation:** John Eoin Healy, Sin Hang Lee.

**Formal Analysis:** Michael John Cook.

**Investigation:** John S. Lambert, John Eoin Healy, Sin Hang Lee.

**Methodology:** Sin Hang Lee.

**Project Administration:** John S. Lambert, Gordana Avramovic.

**Resources:** John S. Lambert, Sin Hang Lee.

**Supervision:** John S. Lambert.

**Validation:** John, S. Lambert.

**Visualization:** John S. Lambert, Ross Murtagh.

**Writing – Original Draft Preparation:** John S. Lambert,

**Writing – Review & Editing:** John S. Lambert, Michael John Cook, John Eoin Healy, Ross Murtagh, Gordana Avramovic, Sin Hang Lee.

